# Brain-age models with lower age prediction accuracy have higher sensitivity for disease detection

**DOI:** 10.1101/2024.03.28.587212

**Authors:** Marc-Andre Schulz, Nys Tjade Siegel, Kerstin Ritter

## Abstract

This study critically reevaluates the utility of brain-age models within the context of detecting neurological and psychiatric disorders, challenging the conventional emphasis on maximizing chronological age prediction accuracy. Our analysis of T1 MRI data from 46,381 UK Biobank participants reveals that simpler machine learning models, and notably those with excessive regularization, demonstrate superior sensitivity to disease-relevant changes compared to their more complex counterparts, despite being less accurate in chronological age prediction. This counterintuitive discovery suggests that models traditionally deemed inferior might, in fact, offer a more meaningful biomarker for brain health by capturing variations pertinent to disease states.

These findings challenge the assumption that accuracy-optimized brain-age models serve as useful normative models of brain aging. Optimizing for age accuracy appears misaligned with normative aims: it drives models to rely on low-variance aging features and to deemphasize higher-variance signals that are more informative about brain health and disease. Consequently, we propose a recalibration of focus towards models that, while less accurate in conventional terms, yield brain-age gaps with larger patient– control effect sizes, offering greater utility in early disease detection and understanding the multifaceted nature of brain aging.

## Introduction

The advancement of neuroimaging techniques has paved the way for innovative measures of brain aging, providing insights into biological aging processes that extend beyond mere chronological metrics (Cole & Franke, 2017). Among these, brain-age prediction models, underpinned by sophisticated machine learning algorithms, have emerged as promising tools for identifying deviations from normative aging trajectories (Baecker et al., 2021). Such deviations, manifesting as discrepancies between predicted and actual brain age, termed the brain-age gap, may signal the early onset of neurological or psychiatric disorders (Chen et al., 2022).

The brain age gap has been used to investigate various diseases of the central nervous system, including Alzheimer’s (Gaser et al., 2013), schizophrenia (Koutsouleris et al., 2014), and major depressive disorder (Han et al., 2021). Evidence suggests that a wider brain-age gap correlates with more severe disease symptoms or faster progression, positioning brain age as a potentially sensitive biomarker for these conditions (Cole et al., 2019).

The pursuit of ever-more precise brain age models, from linear regression to convolutional neural networks to cutting-edge vision transformers, raises critical questions about the relationship between model accuracy and clinical utility. Initial observations suggest that less optimized models might paradoxically reveal more pronounced brain-age gaps, challenging the assumption that increased accuracy inherently enhances clinical value (Bashyam et al., 2020; Hahn et al., 2021; Jirsaraie et al., 2023). Furthermore, recent work by Vidal-Piñeiro et al. (2022) has demonstrated that cross-sectional brain age estimates may primarily reflect stable individual differences rather than ongoing aging processes or pathological changes. This finding calls into question the interpretation of brain-age gaps as direct measures of accelerated aging or disease-related change.

In light of these emerging complexities and apparent contradictions in the field of brain-age modeling, there is a clear need for a comprehensive reevaluation of our approach to developing and applying these models in clinical contexts. This study aims to: (1) demonstrate that simpler and over-regularized models can be more effective biomarkers for detecting neurological and psychiatric disorders, despite lower age prediction accuracy; (2) challenge the conventional focus on maximizing age prediction accuracy in brain-age modeling; and (3) explore the mechanisms underlying the superior performance of simpler models in detecting disease-related brain changes. By ‘better’ or ‘more effective’, we specifically mean generating brain-age gaps that are more sensitive to various conditions, as quantified by larger effect sizes in group comparisons between patients and matched controls.

## Results

Our investigation centers on discerning under which conditions brain age models generate clinically useful brain age gaps. This study analyzed T1 MRI images from 46,381 UK Biobank participants (Sudlow et al., 2015). We focused on a spectrum of psychiatric disorders - alcohol dependency, bipolar disorder, depression - and neurological disorders, such as Parkinson’s disease, epilepsy, and sleep disorders, alongside cognitive and environmental factors including fluid intelligence, recent severe stress exposure, and levels of social support. To ensure rigorous comparison, we applied propensity score matching to generate matched healthy cohorts (Caliendo & Kopeinig, 2008), excluding individuals with any diagnosed neurological or psychiatric condition. Matching criteria included sex, age, education level, household income, the Townsend deprivation index, and genetic principal components, serving as an ethnicity proxy. Fig 1 illustrates our analysis workflow. Detailed information on targets and covariates are provided in the Methods section.

**Fig 1:**
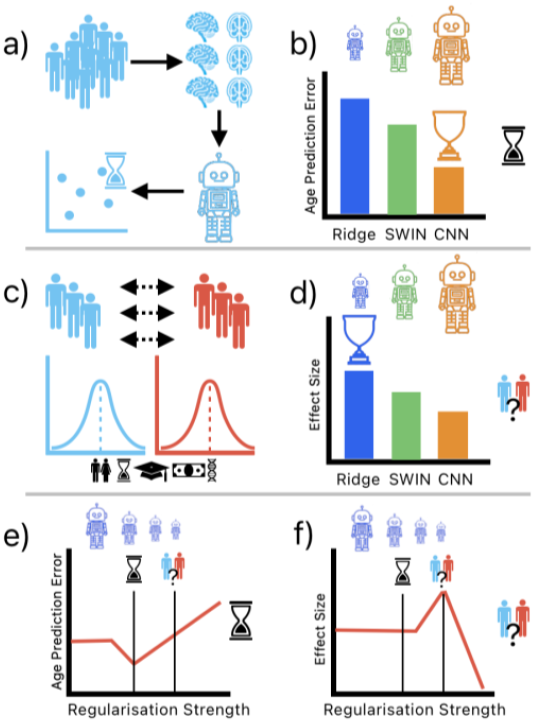
Illustrative Overview of Brain Age Prediction Model Development and Analysis. a) Development of brain age prediction models using T1 brain images from a healthy control group. b) Comparison of model complexity and accuracy, showing Resnet-50 CNN and SWIN vision transformer models surpassing a regularized linear model in age estimation precision. c) Creation of matched patient and healthy control cohorts to examine brain age disparities across various neurological and psychiatric conditions. d) Inverse relationship observed between the machine learning models’ prediction accuracy and the magnitude of brain age gap effect sizes in comparing patient and control groups. e-f) Exploration of the balance between model regularization strength, age prediction accuracy, and the ability to discriminate between patient and control groups, indicating that over-regularization, while reducing age prediction accuracy, enhances patient-control discrimination.

We trained three distinct models to explore these dynamics: a Ridge regularized linear regression model (Hastie et al., 2009), a 3D ResNet-50 CNN (Fisch et al., 2021; He et al., 2016; Kolbeinsson et al., 2020; Siegel et al., 2025), and a 3D SWIN transformer (Hatamizadeh et al., 2022; Liu et al., 2021; Siegel et al., 2025), each chosen for its unique approach to learning from imaging data. The Ridge regression model was selected for its ability to handle high-dimensional data with built-in regularization, making it well-suited for the imaging-derived phenotypes (IDPs; (Alfaro-Almagro et al., 2018). The ResNet-50 CNN was chosen for its proven effectiveness in image analysis tasks, particularly its ability to learn hierarchical features from raw image data. The SWIN transformer was included as a state-of-the-art model capable of capturing long-range dependencies in image data, which may be particularly relevant for whole-brain analysis. The linear model leveraged precomputed imaging-derived phenotypes, while the deep learning models utilized raw, minimally processed T1 images, aiming to capture the most granular features possible.

### Increased Model Accuracy May Diminish Biomarker Effectiveness

We examined how varying degrees of complexity and fine-tuning of brain age models influence their ability to detect brain-age gaps associated with selected psychiatric and neurological conditions. We posited that simpler models, despite their potential limitations in capturing complex patterns, might exhibit heightened sensitivity to variations relevant to disease states. Conversely, more expressive models could potentially mask these signals by fitting to fine-grained but disease invariant aging features.

Our findings corroborate and extend previous observations on the relationship between model complexity and biomarker effectiveness: the simplest model, based on Ridge regression (1.4k parameters), demonstrated the largest brain-age gap effect sizes. Effect sizes appeared to diminish for the more complex and more accurate SWIN transformer (10.1M trainable parameters) and CNN (46.2M trainable parameters) models (see Fig 2). To further investigate this phenomenon, we conducted additional analyses varying key parameters of the model training process. These analyses, presented in Fig 3, explore how effect sizes change under varying training set sizes, different random seeds for model initialization, and increasing numbers of training epochs. The trend of inverse relationships between age prediction accuracy and disease detection sensitivity appears to hold across varying conditions. In sum, we find that conventional methods of improving accuracy (e.g., more expressive models, increasing training data or training duration) can actually decrease the model’s sensitivity to disease-related brain changes.

**Fig 2:**
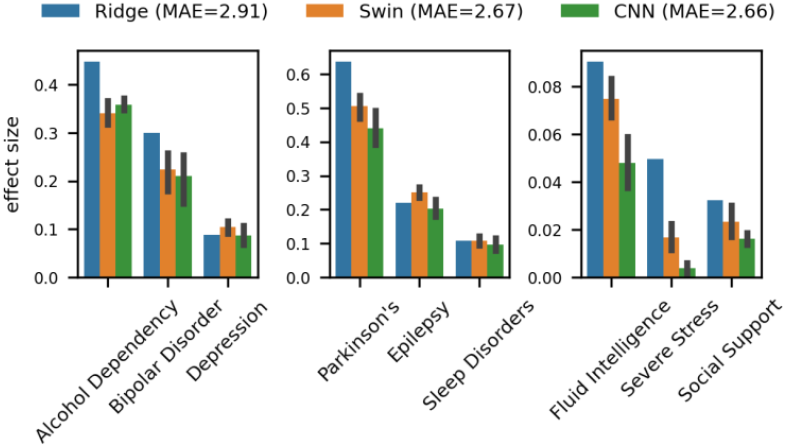
Visualization of Model Complexity vs. Sensitivity in Detecting Brain-Age Gaps. This figure illustrates the nuanced relationship between the complexity of machine learning models and their ability to detect brain-age gaps associated with psychiatric and neurological conditions. It highlights a key observation: models with lower complexity, such as Ridge regression, exhibited a heightened sensitivity in identifying deviations that are clinically significant, compared to their more sophisticated counterparts like CNNs and SWIN Transformers. This unexpected finding challenges conventional expectations, suggesting that simplicity in model architecture may enhance biomarker efficacy by preserving sensitivity to critical, disease-relevant variations. Errorbars indicate standard deviation (STD) over multiple model train runs (with exception of deterministic Ridge model). The data underlying this figure can be found in fig2_data.csv.

**Fig 3:**
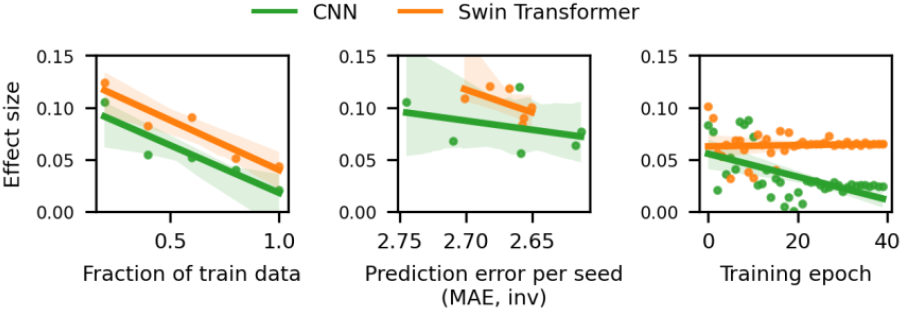
More accurate models can yield worse biomarker effect size. Conventional ways of improving model accuracy like increasing the amount of train data (left), choosing the best performing model out of multiple randomly initialized train runs (middle), and increasing training duration (right) negatively impacted brain age gap effect size for depression for both CNN and SWIN models. We focus on depression as it represents our largest clinical sample, offering the most robust statistical foundation. Similar patterns were observed for a majority of other conditions (see Fig F in S1 Text). Shaded area represents the confidence interval for the regression estimate. The data underlying this figure can be found in fig3_data.csv.

This counterintuitive result underscores a crucial insight: simpler models may possess a greater ability to function as effective biomarkers, hinting that the expressive capacity of complex models exacerbates the impact of an optimization objective that is not fully aligned with normative modelling.

### Over-Regularization of ML Models Increases Biomarker Sensitivity

We investigated how increasing regularization in Ridge regression models affects their ability to generate clinically useful brain-age gaps, hypothesizing that excessive regularization might decrease age prediction accuracy but enhance the model’s utility as a biomarker by improving its sensitivity to disease-relevant anomalies.

We systematically adjusted the L2 regularization strength in a sequence of Ridge regression models tasked with brain age prediction, intentionally exceeding the regularization level that typically optimizes age prediction accuracy (Fig 4). The findings confirmed that while over-regularization reduced the models’ age estimation precision, it notably increased their sensitivity to the brain-age gap associated with selected psychiatric and neurological conditions.

**Fig 4:**
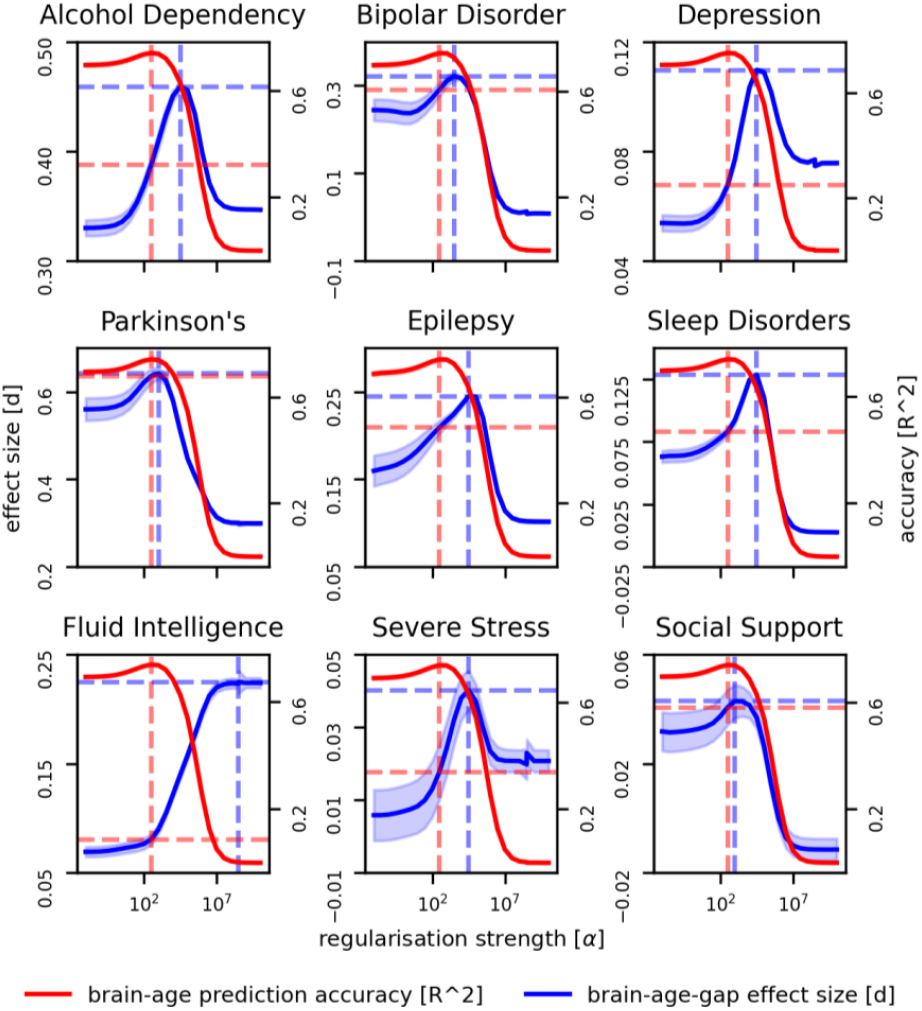
Impact of L2 Regularization on Brain-Age Gap Detection Sensitivity. This figure depicts how adjustments in L2 regularization levels within Ridge regression models influence their ability to identify brain-age gaps associated with selected psychiatric and neurological conditions (effect size, left y-axis, blue) and their age prediction accuracy (mean absolute error, right y-axis, red). The x-axis represents increasing regularization strength (α). The vertical dashed lines demarcate three regions: optimal age prediction (dashed red), transition (middle), and optimal disease detection (dashed blue). In the transition region, we observe an inverse relationship between prediction accuracy and disease detection sensitivity. Horizontal dashed lines indicate effect sizes at α values maximizing prediction accuracy (red) and effect size (blue), i.e. the potential sensitivity gain when prioritizing effect size over prediction accuracy. Patterns vary across conditions, reflecting differences in associated brain changes. Shaded regions indicate standard error derived from bootstrap resampling. The data underlying this figure can be found in fig4_data.csv.

Our analysis revealed three distinct regions of regularization strength: optimal age prediction, transition, and optimal disease detection. In the transition region, we observed an inverse relationship between prediction accuracy and disease detection sensitivity across all conditions. As regularization increased, age prediction error rose while the effect size for detecting condition-related brain-age gaps improved. The regularization strength that optimized age prediction accuracy (typically at lower α values) differed from the one that maximized discriminative ability (often at higher α values). This difference represents a potential gain in biomarker sensitivity when prioritizing effect size over prediction accuracy.

Interestingly, the effect size often peaks at an intermediate level of regularization, rather than at the maximum level. This pattern suggests an optimal balance between model simplicity and feature retention. This balance point varies across conditions, potentially reflecting differences in the nature and extent of associated brain changes.

The results suggest that increasing the level of regularization, effectively simplifying the model’s representation of aging patterns, enhances its ability to identify variations pertinent to clinical conditions, distinct from conventional aging. This loss in precision for age prediction, counterintuitively, did not diminish the model’s value; rather, it bolstered its performance as a biomarker. Such findings support the notion that model complexity negatively affects how well the brain-age gap encompasses deviations from a normative aging process, affirming the potential of strategically simplified models to serve as more effective tools in disease detection and characterization.

### Overregularized models focus on global gray matter volume

To gain insight into the mechanisms underlying the improved biomarker performance of over-regularized models, we analyzed the feature importances of Ridge regression models with varying levels of regularization. We employed SHapley Additive exPlanations (SHAP) values to quantify the contribution of each feature to the model’s predictions (Lundberg & Lee, 2017).

Our analysis revealed a clear trend: as regularization increased, the models increasingly relied on global measures of brain structure, particularly total gray matter volume. This shift in focus from localized features to global indicators aligns with our understanding of many neurological and psychiatric conditions, which often manifest as widespread alterations in brain structure rather than changes confined to specific regions (Pini et al., 2016; Schmaal et al., 2017; Van Erp et al., 2018).

When comparing the top features for both accuracy-optimized and disease-detection-optimized models, we observed distinct patterns of feature importance. The accuracy-optimized model prioritizes features such as specific regional volumes and contrasts, which are strong predictors of age (Raz et al., 2010). In contrast, the disease-detection-optimized model focuses more on global measures like total gray matter volume and ventricular size, which may be more sensitive to a range of pathological changes. These differences are visually represented in Fig 5.

**Fig 5:**
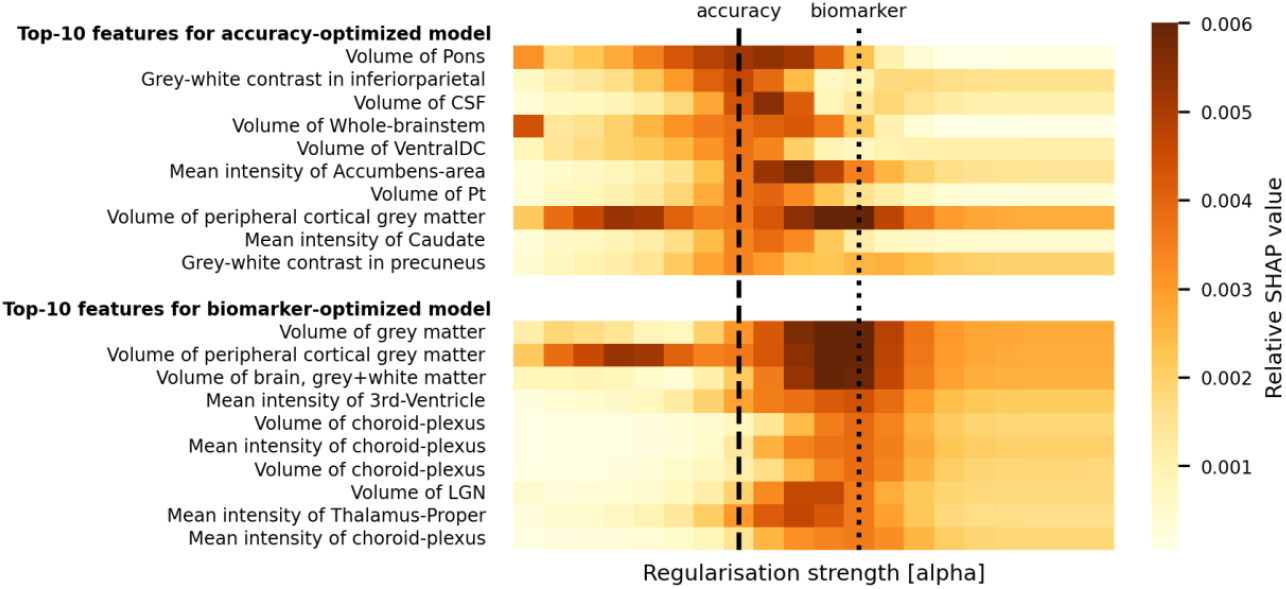
Feature importances for brain age prediction using Ridge regression. The heatmaps illustrate the top-10 features for both the accuracy-optimized model (top panel) and the biomarker-optimized model (bottom panel), with color intensity representing the relative SHAP values. The x-axis denotes increasing regularization strength [alpha]. The accuracy-optimized model (α≈10^3), trained for maximum age prediction accuracy, highlights features such as the volume of the pons, grey-white contrast in the inferior parietal region, and volume of the cerebrospinal fluid (CSF). Conversely, the biomarker-optimized model (α≈10^5), trained to enhance the brain age gap effect size for a majority of conditions (cf. Methods and Fig E in S1 Text) versus controls, emphasizes features like the volume of grey matter, volume of peripheral cortical grey matter, and mean intensity of the third ventricle. The dashed vertical lines indicate the regularization strength [alpha] at which each model was optimized. This comparison underscores the distinct feature importances between models focused on accuracy versus those optimized for sensitivity to disease-relevant changes, supporting the manuscript’s thesis that traditional accuracy-optimized models may not provide the best biomarkers for disease detection. The data underlying this figure can be found in fig5_data.csv.

This finding offers a plausible explanation for the superior performance of over-regularized models as biomarkers. By focusing on global brain characteristics, these models may be more sensitive to the diffuse structural changes associated with various brain disorders. Conversely, more complex models that can capture intricate, localized patterns may paradoxically be less effective at detecting these broad, disease-related alterations.

Our interpretability analysis thus provides mechanistic insights into why simpler, over-regularized models can outperform more complex ones in disease detection. By prioritizing global brain measures, these models appear to capture broader indicators of brain health that are particularly relevant to a range of neurological and psychiatric conditions. This underscores the importance of aligning model complexity with the intended application, suggesting that in the context of disease detection, strategically simplified models may offer advantages over those optimized solely for age prediction accuracy.

## Discussion

Our study reassesses the efficacy of brain-age models in detecting neurological and psychiatric disorders, revealing that simpler or more regularized models often demonstrate enhanced sensitivity to disease-related variations compared to more complex counterparts. This finding challenges the conventional assumption that higher age prediction accuracy necessarily leads to better biomarkers, suggesting a need to reevaluate criteria for assessing brain-age models in clinical and research settings.

The inverse relationship between age prediction accuracy and disease detection sensitivity persisted across various experimental conditions, including different training set sizes, random initializations, and training durations. Our results align with and extend previous work (Bashyam et al., 2020; Jirsaraie et al., 2023), providing systematic evidence across a broader range of models and conditions.

These dynamics help reconcile prior mixed findings about what brain-age reflects (Anatürk et al., 2021; Cole et al., 2019; Nguyen et al., 2024; Vidal-Piñeiro et al., 2021; Wrigglesworth et al., 2022): simpler or more constrained models produce larger patient-control effect sizes, indicating that the brain-age gap integrates both age-normative and pathology-sensitive variation, with the balance governed by training choices.

This work also resonates with recent studies questioning the universal superiority of deep learning over simpler linear models in brain imaging analyses (Schulz, 2020, 2024; Han, 2022; He, 2020). Our results provide further empirical support for the potential advantages of simpler or deliberately constrained models in certain neuroimaging tasks.

Further, our findings align with a methodological shift in the parallel field of epigenetic clocks, where clinical utility has been enhanced by moving beyond the singular goal of predicting chronological age. An analogue to our ‘brain-age paradox’ was demonstrated by Zhang et al. (2019), who found that as an epigenetic clock’s age-prediction accuracy improved, its association with mortality risk attenuated. These convergent findings are reinforced by evidence showing that ‘next-generation’ epigenetic clocks like DunedinPACE and GrimAge, which are less correlated with age, consistently outperform their predecessors in predicting clinical outcomes, including dementia (Sugden et al., 2022), adverse brain structure (Whitman et al., 2024), and frailty and mortality in vulnerable populations (Guida et al., 2024). Furthermore, these advanced clocks show greater sensitivity to socioeconomic and lifestyle risk factors even in younger adults (Harris et al., 2024). Our work suggests that simpler, over-regularized brain-age models function analogously, de-emphasizing pure age prediction to better capture the clinically relevant signals of pathology.

The preference for simpler or more strongly regularized models as biomarkers can be explained by how the optimization objective interacts with population variance. In probabilistic normative modeling, one estimates the distribution of selected features conditional on covariates (e.g., age, sex, site) and then quantifies each individual’s deviation from that distribution. This paradigm relies on leveraging cross-sectional variation in features that are sensitive to brain health and disease. Brain-age models, by contrast, compress many features into a single scalar trained to minimize chronological age error. To reduce average prediction error, they place more weight on features with high signal-to-noise for age and low residual variability across individuals. As a result, features with greater cross-sectional variability, often those most sensitive to pathology, are down-weighted or ignored. This can yield excellent age-prediction accuracy while producing brain-age gaps that are less informative for detecting disease-related deviations. In short, optimization for age accuracy can attenuate precisely the variance that normative modeling aims to characterize.

This misalignment can be amplified by cohort contamination. If the “healthy” training set includes undiagnosed cases or uses broad inclusion criteria, disease-linked variation appears as noise with respect to age. Because the training procedure seeks features that predict age consistently across the sample, the model will preferentially rely on disease-invariant aging signals over features whose age trajectory is perturbed by illness, further reducing sensitivity to pathology at test time. Complete elimination of such contamination is challenging in large population cohorts, as many disorders manifest as tail deviations on otherwise continuous neurobiological dimensions.

Further, regularization encourages a focus on robust features, potentially corresponding to larger-scale structural changes more likely associated with disease processes. This suggests an optimal balance point in the bias-variance trade-off that differs for disease detection versus age prediction, where moderate regularization constraints maintain sensitivity to disease-relevant patterns while avoiding fitting to fine-grained age-specific features.. Simpler models might exhibit greater robustness to the heterogeneity of brain changes in disease, capturing general indicators of abnormality across diverse presentations. Collectively, these attributes suggest that the constraints imposed by simpler or more regularized models may inadvertently align with the nature of the biological signals most relevant to disease detection, explaining their enhanced performance as biomarkers despite lower age prediction accuracy.

Our findings provide empirical support for these theoretical advantages. Feature importance analysis (Fig 4) reveals that as regularization increases in the Ridge regression model, it tends to rely more heavily on global measures of brain structure, particularly total gray matter volume. This shift towards global features aligns well with the widespread alterations often seen in neurological and psychiatric conditions (Pini et al., 2016; Schmaal et al., 2017; Van Erp et al., 2018). While our feature importance analysis demonstrates this for linear models, the exact mechanisms in non-linear models remain to be explored. More complex models might learn to prioritize fine-grained, localized patterns that excel in age prediction but are less sensitive to diffuse pathological changes, though directly testing this hypothesis requires further research.

It’s important to note that our findings, while reminiscent of the ‘loose fitting’ hypothesis proposed by Bashyam et al. (2020), differ in crucial ways that address the concerns raised by Hahn (2021). Unlike ‘loose fitting’, or underfitting, which potentially violates normative modeling principles, our approach focuses on model simplicity and regularization rather than intentional undertraining. Undertraining deep networks can yield partially random feature spaces that may coincidentally capture disease-relevant patterns, similar to how randomly initialized CNNs can sometimes provide useful embeddings. In contrast, our regularized models are fully trained but deliberately constrained in their capacity to combine age-relevant features. This means they still learn age-specific patterns, but are limited in how many features they can utilize or how complexly they can combine them. The systematic relationship we observe between regularization strength and biomarker sensitivity suggests that this controlled constraint of age-related feature combinations, rather than random or incomplete feature learning, drives the improved disease detection.

Our study used both minimally processed T1 images and imaging-derived phenotypes. While raw images retain more information, they require significant computational resources and engineering expertise. IDPs offer reduced computational demands, easier interpretability, and lower expertise requirements, but may lose some fine-grained information. Notably, our IDP-based model outperformed those using minimally processed images in detecting disease-related changes, suggesting that pre-extracted features may be sufficient and even advantageous for identifying broad, disease-related patterns.

This highlights a key trade-off in brain-age modeling between information retention and feature robustness. Our results indicate that for developing sensitive biomarkers, IDPs with simpler, more regularized models may offer a good balance of performance, interpretability, and accessibility.

However, this study is not without its limitations. Our analysis was confined to a limited set of target phenotypes and utilized a limited range of machine learning models. Future research should aim to replicate these findings across a more diverse array of diseases and model architectures to validate the generalizability of our conclusions. Additionally, the mechanisms by which simpler or more regularized models achieve greater sensitivity to pathological signals warrant further investigation. Disentangling the features and patterns these models prioritize could offer valuable insights into the biological underpinnings of the brain-age gap phenomenon.

Furthermore, the time elapsed between diagnosis and brain imaging varies across conditions and individuals, potentially affecting the observed effect sizes differently for progressive versus stable conditions. While fluid intelligence showed notably different patterns compared to clinical conditions, the underlying mechanisms for these differences remain to be explored. Future work should investigate how the temporal dynamics of different conditions interact with brain age predictions

Future work could also explore whether systematic regularization of non-linear deep learning models yields similar or potentially even stronger biomarker properties, though this presents significant computational challenges given the high dimensionality of these models’ parameter spaces.

In conclusion, our work contributes to a growing body of evidence that questions the exclusive focus on chronological age prediction accuracy in brain-age modeling. By highlighting the potential of simpler and more regularized models to serve as effective biomarkers for neurological and psychiatric disorders, we advocate for a shift towards models that prioritize clinical relevance and interpretability. This paradigm shift could pave the way for more effective diagnostic tools and intervention strategies, ultimately enhancing patient care in the realm of neurology and psychiatry.

## Methods

### Dataset and Preprocessing

The study utilized T1-weighted magnetic resonance imaging data from 46,381 participants enrolled in the UK Biobank project (Sudlow et al., 2015). This large-scale biomedical database and research resource contains in-depth genetic and health information from half a million UK participants, with a subset undergoing extensive brain imaging. Our analysis used T1 imaging-derived-phenotypes (IDPs; regional gray and white matter volumes, cortical thickness and surface area) provided by the UK Biobank (Alfaro-Almagro et al., 2018), specifically 1425 descriptors defined in UK Biobank category 110. Psychiatric disorders (alcohol dependency, bipolar disorder, depression) and neurology-related disorders (Parkinson’s disease, epilepsy, sleep disorders), as well as cognitive and environmental factors (fluid intelligence, exposure to severe stress in the last 2 years, level of social support) served as prediction targets. Disorders refer to ICD-10 codes F10, F31, F32, G20, G40, and G47, with time of first diagnosis preceding the date of imaging. Fluid intelligence refers to UKB-field 20016, binarized into top and bottom quartile. Exposure to severe stress refers to UKB-field 6145, binarized into having or not having an adverse life event in the last 2 years. Level of social support, specifically “able to confide”, field UKB-2110, was binarized into the majority category almost daily versus anything less frequent. For model training, we excluded all participants who had received any neurological or behavioral (ICD-10 category F or G) diagnosis at any point during follow-up, including diagnoses made after imaging. For the analysis of disease effects, we considered participants “healthy controls” if they had no relevant diagnosis at the time of MRI measurement. For non-disease targets (fluid intelligence, stress, social support), we relaxed this strict comorbidity exclusion to maximize available sample sizes. All features were standard scaled, participants with missing data were excluded on an analysis-by-analysis basis. For details on methodological choices please refer to Note 1 in S1 Text.

### Matching

To investigate the association between brain age gaps and various psychiatric and neurological conditions, we employed propensity score matching (Caliendo & Kopeinig, 2008), with a caliper of 0.25 standard deviations. This technique allowed us to create balanced cohorts of participants with and without specific diagnoses, thereby reducing confounding factors. Matching criteria included sex (field 31), age (field 21003), years of education (field 6138, translated into years of education via the 1997 International Standard Classification of Education), household income (field 738), Townsend deprivation index (field 22189), and the first three genetic principal components (field 22009) as proxies for ethnicity. The resulting sample sizes after matching were: alcohol dependency (n=495), bipolar disorder (n=126), depression (n=1146), Parkinson’s disease (n=95), epilepsy (n=327), sleep disorders (n=1007), fluid intelligence (n=3285), severe stress (n=6528), and social support (n=4250). Healthy controls were drawn from the pool of participants that were not used for training in the respective experiments.

### Brain Age Gap Calculation

Brain age gaps were calculated by training a machine learning model to predict the participants age from the T1 image on a healthy (see sections below) cohort, then using this model to predict held-out participants’ ages. The difference between predicted age and true age was linear sample-level bias corrected (Smith et al., 2019), i.e. the difference between predicted age and true age was bias-corrected using linear regression on all test data, with subsequent analyses performed on the residuals - referred to as the participant’s (corrected) “brain age gap” in our analyses. Throughout this study, we distinguish between the brain-age gap itself (the difference between predicted and chronological age) and the ability of these gaps to discriminate between patient and control groups (quantified by effect sizes).

### Effect Size Calculation

As a proxy for the practical usefulness of a brain age model we calculated effect sizes for our binary target variables (disease status, binarized fluid intelligence, etc.; see above). Specifically, we created a matched control group for each variable via propensity score matching (see above), calculated the brain age gaps for participants in both groups, then used these brain age gaps to calculate the effect size (Cohen’s d) of the given group comparison.

### Experiment 1: Model Training and Architecture Specification

Three distinct machine learning models were trained for the task of brain age prediction: a regularized linear regression model (Hastie et al., 2001), a 3D ResNet-50 convolutional neural network (Fisch et al., 2021; He et al., 2016; Kolbeinsson et al., 2020; Siegel et al. 2025), and a 3D SWIN transformer (Hatamizadeh et al., 2022; Liu et al., 2021; Siegel et al. 2025). The linear model utilized precomputed IDPs, while the deep learning models were trained on minimally processed (skull stripped and linearly registered, mask and transformation matrix provided by UK Biobank) T1 images. Data was split into healthy participants (no neurological or behavioral, i.e. ICD-10 category F or G diagnoses) and patients (any ICD-10 F or G diagnoses). Patients (n=17671) and 1000 random healthy participants were kept as a held-out test set, patients with diagnoses that were not investigated in this work were discarded, and the remaining healthy participants constituted the train set (n=27,538). Brain age prediction accuracy was computed on the healthy test set participants only.

Architecture details and training hyperparameters of CNN and transformer are described in detail in (Siegel et al. 2025). Both deep architectures were trained using AdamW, optimizing the mean squared error loss, using a one-cycle learning rate policy with a maximal learning rate of 10^-2 for the ResNet and 10^-4 for the SWIN transformer. Both were trained for 150,000 gradient update steps, with an effective batch size of 8. Each architecture was trained 6 times with different random initializations. We generally report end-of-training performance and, unless differently specified, the average performance over re-trained instances.

The linear model relied on the ScikitLearn (Pedregosa et al., 2011) RidgeCV implementation (alpha 10^-5 to 10^5, 100 ticks, log spaced).

### Experiment 2: Over-Regularization and Biomarker Sensitivity

We systematically varied the regularization strength in the Ridge regression brain age model to study its impact on age prediction accuracy and the detection of brain-age gaps. This experiment aimed to understand how over-regularization, or intentionally reducing model complexity, might influence the model’s utility as a biomarker for disease. The train set size was reduced to 5000 to ensure that both over-regularization and under-regularization effects could be visualized and easily distinguished. The train set was randomly sampled 10 times, so that values in Fig 3 refer to the mean over brain age models that were trained on different sets of participants, and error bars refer to the standard error (SE) respectively.

### Experiment 3: Interpretability Analysis

To further elucidate why simpler, over-regularized models are more effective in detecting disease-relevant changes, we conducted an interpretability experiment. This experiment compares the feature importances of Ridge models with increasing regularization strength. Models were trained in the same manner as in Experiment 2. Feature importances were derived using SHapley Additive exPlanations (SHAP), averaged over the 10 random train set resamplings. In Fig 4 we present the Top-10 important features for the accuracy-maximizing regularization strength and the regularization strength maximizing effect sizes for a majority of conditions (alcohol dependence, depression, epilepsy, sleep disorders, stress). Top-10 important features for higher level of regularization are presented in Fig E in S1 Text.

### Statistical Analysis Details

Estimation of statistical uncertainties was performed via repeated random sub-sampling validation; reported uncertainties refer to variability over models trained on different subsets of the data. Mean Absolute Error served as our primary metric for assessing age prediction accuracy, while Cohen’s d was used to quantify the effect sizes in a manner that is both standardized and interpretable. Analyses were conducted using Python 3.10.8 with the following key library versions: scikit-learn 1.3.0, PyTorch 1.12.1, numpy 1.25.2, and scipy 1.11.1. Deep learning models were trained using NVIDIA A100 GPUs with CUDA 12.2.

### Ethical Considerations

This study was conducted under the umbrella of the UK Biobank’s ethics agreements. The UK Biobank has received ethical approval from the National Health Service National Research Ethics Service (16/NW/0274) to collect and distribute data to approved researchers compliant with the declaration of Helsinki. As our study utilized de-identified data provided by the UK Biobank, individual consent from participants was covered under the UK Biobank’s broad consent model.

## Acknowledgments

We thank Roshan Rane, Habakuk Hain, Ruben Brandhofer, and Marcel Jühling for feedback on the manuscript. We thank the UKBB participants for their voluntary commitment and the UKBB team for their work in collecting, processing, and disseminating these data for analysis. Research was conducted using the UKBB resource under project-ID 33073. The authors acknowledge the Scientific Computing of the IT Division at the *Charité - Universitätsmedizin Berlin* for providing computational resources that have contributed to the research results reported in this paper, as well as support from Gemeinnützige Hertie Stiftung.

## Supplementary Figures

S1 Text: Supplementary Online Material

## References

Alfaro-Almagro, F., Jenkinson, M., Bangerter, N. K., Andersson, J. L. R., Griffanti, L., Douaud, G., Sotiropoulos, S. N., Jbabdi, S., Hernandez-Fernandez, M., Vallee, E., Vidaurre, D., Webster, M., McCarthy, P., Rorden, C., Daducci, A., Alexander, D. C., Zhang, H., Dragonu, I., Matthews, P. M., … Smith, S. M. (2018). Image processing and Quality Control for the first 10,000 brain imaging datasets from UK Biobank. NeuroImage, 166, 400–424.

Baecker, L., Garcia-Dias, R., Vieira, S., Scarpazza, C., & Mechelli, A. (2021). Machine learning for brain age prediction: Introduction to methods and clinical applications. EBioMedicine, 72, 103600.

Caliendo, M., & Kopeinig, S. (2008). Some practical guidance for the implementation of propensity score matching. Journal of Economic Surveys, 22(1), 31–72.

Chen, C.-L., Hwang, T.-J., Tung, Y.-H., Yang, L.-Y., Hsu, Y.-C., Liu, C.-M., Lin, Y.-T., Hsieh, M.-H., Liu, C.-C., Chien, Y.-L., Hwu, H.-G., & Tseng, W.-Y. I. (2022). Detection of advanced brain aging in schizophrenia and its structural underpinning by using normative brain age metrics. NeuroImage. Clinical, 34, 103003.

Cole, J. H., & Franke, K. (2017). Predicting Age Using Neuroimaging: Innovative Brain Ageing Biomarkers. Trends in Neurosciences, 40(12), 681–690.

Cole, J. H., Marioni, R. E., Harris, S. E., & Deary, I. J. (2019). Brain age and other bodily “ages”: implications for neuropsychiatry. Molecular Psychiatry, 24(2), 266–281.

Fisch, L., Ernsting, J., Winter, N. R., Holstein, V., Leenings, R., Beisemann, M., Sarink, K., Emden, D., Opel, N., Redlich, R., Repple, J., Grotegerd, D., Meinert, S., Wulms, N., Minnerup, H., Hirsch, J. G., Niendorf, T., Endemann, B., Bamberg, F., … Hahn, T. (2021). Predicting brain-age from raw T 1 - weighted Magnetic Resonance Imaging data using 3D Convolutional Neural Networks. In arXiv [eess.IV]. http://arxiv.org/abs/2103.11695

Gaser, C., Franke, K., Klöppel, S., Koutsouleris, N., Sauer, H., & Alzheimer’s Disease Neuroimaging Initiative. (2013). BrainAGE in Mild Cognitive Impaired Patients: Predicting the Conversion to Alzheimer’s Disease. PloS One, 8(6), e67346.

Han, L. K. M., Dinga, R., Hahn, T., Ching, C. R. K., Eyler, L. T., Aftanas, L., Aghajani, M., Aleman, A., Baune, B. T., Berger, K., Brak, I., Filho, G. B., Carballedo, A., Connolly, C. G., Couvy-Duchesne, B., Cullen, K. R., Dannlowski, U., Davey, C. G., Dima, D., … Schmaal, L. (2021). Brain aging in major depressive disorder: results from the ENIGMA major depressive disorder working group. Molecular Psychiatry, 26(9), 5124–5139.

Hastie, T., Tibshirani, R., & Friedman, J. (2009). The Elements of Statistical Learning: Data Mining, Inference, and Prediction. Springer, New York, NY.

Hastie, T., Tibshirani, T., & Friedman, J. (2001). The elements of statistical learning (Vol. 1). Springer series in statistics New York.

Hatamizadeh, A., Nath, V., Tang, Y., Yang, D., Roth, H. R., & Xu, D. (2022). Swin UNETR: Swin Transformers for Semantic Segmentation of Brain Tumors in MRI Images. Brainlesion: Glioma, Multiple Sclerosis, Stroke and Traumatic Brain Injuries, 272–284.

He, K., Zhang, X., Ren, S., & Sun, J. (2016). Deep residual learning for image recognition. Proceedings of the IEEE Conference on Computer Vision and Pattern Recognition,770–778.

Kolbeinsson, A., Filippi, S., Panagakis, Y., Matthews, P. M., Elliott, P., Dehghan, A., & Tzoulaki, I. (2020). Accelerated MRI-predicted brain ageing and its associations with cardiometabolic and brain disorders. Scientific Reports, 10(1), 19940.

Koutsouleris, N., Davatzikos, C., Borgwardt, S., Gaser, C., Bottlender, R., Frodl, T., Falkai, P., Riecher-Rössler, A., Möller, H.-J., Reiser, M., Pantelis, C., & Meisenzahl, E. (2014). Accelerated brain aging in schizophrenia and beyond: a neuroanatomical marker of psychiatric disorders. Schizophrenia Bulletin, 40(5), 1140–1153.

Liu, Z., Lin, Y., Cao, Y., Hu, H., Wei, Y., Zhang, Z., Lin, S., & Guo, B. (2021). Swin Transformer: Hierarchical Vision Transformer using Shifted Windows. In arXiv [cs.CV]. http://arxiv.org/abs/2103.14030

Pedregosa, F., Varoquaux, G., Gramfort, A., Michel, V., Thirion, B., Grisel, O., Blondel, M., Prettenhofer, P., Weiss, R., Dubourg, V., Vanderplas, J., Passos, A., Cournapeau, D., Brucher, M., Perrot, M., & Duchesnay, É. (2011). Scikit-learn: Machine Learning in Python. Journal of Machine Learning Research: JMLR, 12(Oct), 2825–2830.

Pini, L., Pievani, M., Bocchetta, M., Altomare, D., Bosco, P., Cavedo, E., Galluzzi, S., Marizzoni, M., & Frisoni, G. B. (2016). Brain atrophy in Alzheimer’s Disease and aging. Ageing Research Reviews, 30, 25–48.

Raz, N., Ghisletta, P., Rodrigue, K. M., Kennedy, K. M., & Lindenberger, U. (2010). Trajectories of brain aging in middle-aged and older adults: regional and individual differences. NeuroImage, 51(2), 501– 511.

Schmaal, L., Hibar, D. P., Sämann, P. G., Hall, G. B., Baune, B. T., Jahanshad, N., Cheung, J. W., van Erp, T. G. M., Bos, D., Ikram, M. A., Vernooij, M. W., Niessen, W. J., Tiemeier, H., Hofman, A., Wittfeld, K., Grabe, H. J., Janowitz, D., Bülow, R., Selonke, M., … Veltman, D. J. (2017). Cortical abnormalities in adults and adolescents with major depression based on brain scans from 20 cohorts worldwide in the ENIGMA Major Depressive Disorder Working Group. Molecular Psychiatry, 22(6), 900–909.

Siegel, N. T., Kainmueller, D., Deniz, F., Ritter, K., & Schulz, M. A. (2025). Do transformers and CNNs learn different concepts of brain age?. Human Brain Mapping, 46(8), e70243.

Smith, S. M., Vidaurre, D., Alfaro-Almagro, F., Nichols, T. E., & Miller, K. L. (2019). Estimation of brain age delta from brain imaging. NeuroImage, 200, 528–539.

Sudlow, C., Gallacher, J., Allen, N., Beral, V., Burton, P., Danesh, J., Downey, P., Elliott, P., Green, J., Landray, M., Liu, B., Matthews, P., Ong, G., Pell, J., Silman, A., Young, A., Sprosen, T., Peakman, T., & Collins, R. (2015). UK biobank: an open access resource for identifying the causes of a wide range of complex diseases of middle and old age. PLoS Medicine, 12(3), e1001779.

Van Erp, T. G. M., Walton, E., Hibar, D. P., Schmaal, L., Jiang, W., Glahn, D. C., Pearlson, G. D., Yao, N., Fukunaga, M., Hashimoto, R., & Others. (2018). Cortical brain abnormalities in 4474 individuals with schizophrenia and 5098 control subjects via the enhancing neuro imaging genetics through meta analysis (ENIGMA) consortium. Biological Psychiatry, 84(9), 644–654.

